# Significant associations between driver gene mutations and DNA methylation alterations across many cancer types

**DOI:** 10.1101/145680

**Authors:** Yun-Ching Chen, Valer Gotea, Gennady Margolin, Laura Elnitski

**Author notes:** Corresponding author (LE).

## Abstract

Recent evidence shows that mutations in several driver genes can cause aberrant methylation patterns, a hallmark of cancer. In light of these findings, we hypothesized that the landscapes of tumor genomes and epigenomes are tightly interconnected. We measured this relationship using principal component analyses and methylation-mutation associations applied at the nucleotide level and with respect to genome-wide trends. We found a few mutated driver genes were associated with genome-wide patterns of aberrant hypomethylation or CpG island hypermethylation in specific cancer types. We identified associations between 737 mutated driver genes and site-specific methylation changes. Moreover, using these mutation-methylation associations, we were able to distinguish between two uterine and two thyroid cancer subtypes. The driver gene mutation-associated methylation differences between the thyroid cancer subtypes were linked to differential gene expression in JAK-STAT signaling, NADPH oxidation, and other cancer-related pathways. These results establish that driver-gene mutations are associated with methylation alterations capable of shaping regulatory network functions. In addition, the methodology presented here can be used to subdivide tumors into more homogeneous subsets corresponding to their underlying molecular characteristics, which could improve treatment efficacy.

**Author summary:** Mutations that alter the function of driver genes by changing DNA nucleotides have been recognized as a key player in cancer progression. Recent evidence showed that DNA methylation, a molecular signature that is used for controlling gene expression and that consists of cytosine residues with attached methyl groups in the context of CG dinucleotides, is also highly dysregulated in cancer and contributes to carcinogenesis. However, whether those methylation alterations correspond to mutated driver genes in cancer remains unclear. In this study, we analyzed 4,302 tumors from 18 cancer types and demonstrated that driver gene mutations are inherently connected with the aberrant DNA methylation landscape in cancer. We showed that those driver gene-associated methylation patterns can classify heterogeneous tumors in a cancer type into homogeneous subtypes and have the potential to influence the genes that contribute to tumor growth. This finding could help us to better understand the fundamental connection between driver gene mutations and DNA methylation alterations in cancer and to further improve the cancer treatment.

## Introduction

DNA methylation (DNAm) is highly dysregulated in cancers from many organs [1, 2] where it is characterized by aberrant CpG island (CGI) hypermethylation and long-range blocks of hypomethylation. Moreover, dysregulated DNAm at specific locations within the genome often distinguishes heterogeneous tumors within cancer types into homogeneous subtypes [3, 4]. The origin of these dramatic changes in the DNAm of tumor cells remains a puzzle. On the one hand, DNAm alterations at particular CpG sites in tumors are associated with the aging process in normal cells [5, 6]. This has led some researchers to propose that cell proliferation, which drives age-associated DNAm errors in normal cells, is also responsible for aberrant DNAm in cancer [7]. On the other hand, although these DNAm errors exhibit a linear association with the number of cell divisions in normal cells, in cancer cells, they are not well correlated with the mRNA expression-based mitotic index in many cancer types [7]. This suggests that in tumor cells, some factor other than cell proliferation is shaping the DNAm landscape. Because tumors of the same molecular subtype often harbor both dysregulated DNAm at particular locations in the genome and mutations in driver genes [3, 8], we decided to investigate more broadly the connection between somatic mutations and specific aberrant DNAm patterns.

Several genes that are known to play a defining role in a wide variety of cancer types [9, 10], such as *TP53, IDH1, BRAF*, and *KRAS*, have been functionally linked to dysregulated DNA methylation. For example, in hepatocellular carcinoma and esophageal squamous cell carcinoma, *TP53* mutations are associated with extensive DNA hypomethylation [11, 12]. In glioblastoma, *IDH1* mutations produce widespread CGI hypermethylation, termed the CpG island methylator phenotype (CIMP), by inhibiting the TET-demethylation pathway [13, 14]. In colorectal cancer, the BRAF V600E mutation results in DNA hypermethylation and CIMP development by upregulating the transcriptional repressor MAFG, which recruits the DNA methyltransferase DNMT3B to its targets at promoter CGIs [15]. Likewise, the KRAS G13D mutation upregulates another transcriptional repressor, ZNF304, to establish a CIMP-intermediate pattern in colorectal cancer [16].

Based on these findings, we hypothesized that the genomic and epigenetic landscapes were stable and interdependent and therefore specific driver mutations would correlate with specific DNAm patterns. Thus, in this study, we systematically evaluated mutation-methylation associations across 4,302 tumors from 18 cancer types, along with 727 normal tissue samples from The Cancer Genome Atlas (TCGA). By investigating DNAm alterations associated with mutated driver genes on both a genome-wide scale and a site-specific scale, we were able to show that i) mutated driver genes are tightly associated with DNAm variation in cancer; ii) some associations are present across cancer types, whereas others are cancer type-specific; iii) mutation-methylation associations cannot be explained by cell proliferation alone; and iv) these associations can be used to classify tumors into molecular subtypes and gain insight into functional alterations. Together, these results establish that driver mutations and DNAm alterations are tightly coupled in tumor cells, and that this coupling may affect important regulatory networks related to oncogenesis.

## Results

### Association between driver gene mutations and methylation patterns in cancer

To determine whether mutated driver genes were associated with methylation changes, first we performed principal component analysis (PCA) on methylation data for each of 18 different cancer types; within a given cancer type, tumor samples were projected onto the principal components (PCs). Illumina Infinium human methylation 450K array data and somatic mutation data were downloaded from TCGA (Table 1), and driver genes were predicted with MutSigCV [17] (see Materials and Methods for details). For each cancer type, a driver gene was considered to be associated with a PC if samples in which the gene was mutated were unevenly distributed toward the positive or negative extremes of that PC (q<0.05; two sided Wilcoxon rank-sum test). We assessed each driver gene for the top five DNAm PCs; examples of PC1-associated driver genes are shown in Fig 1A.

**Fig 1.**
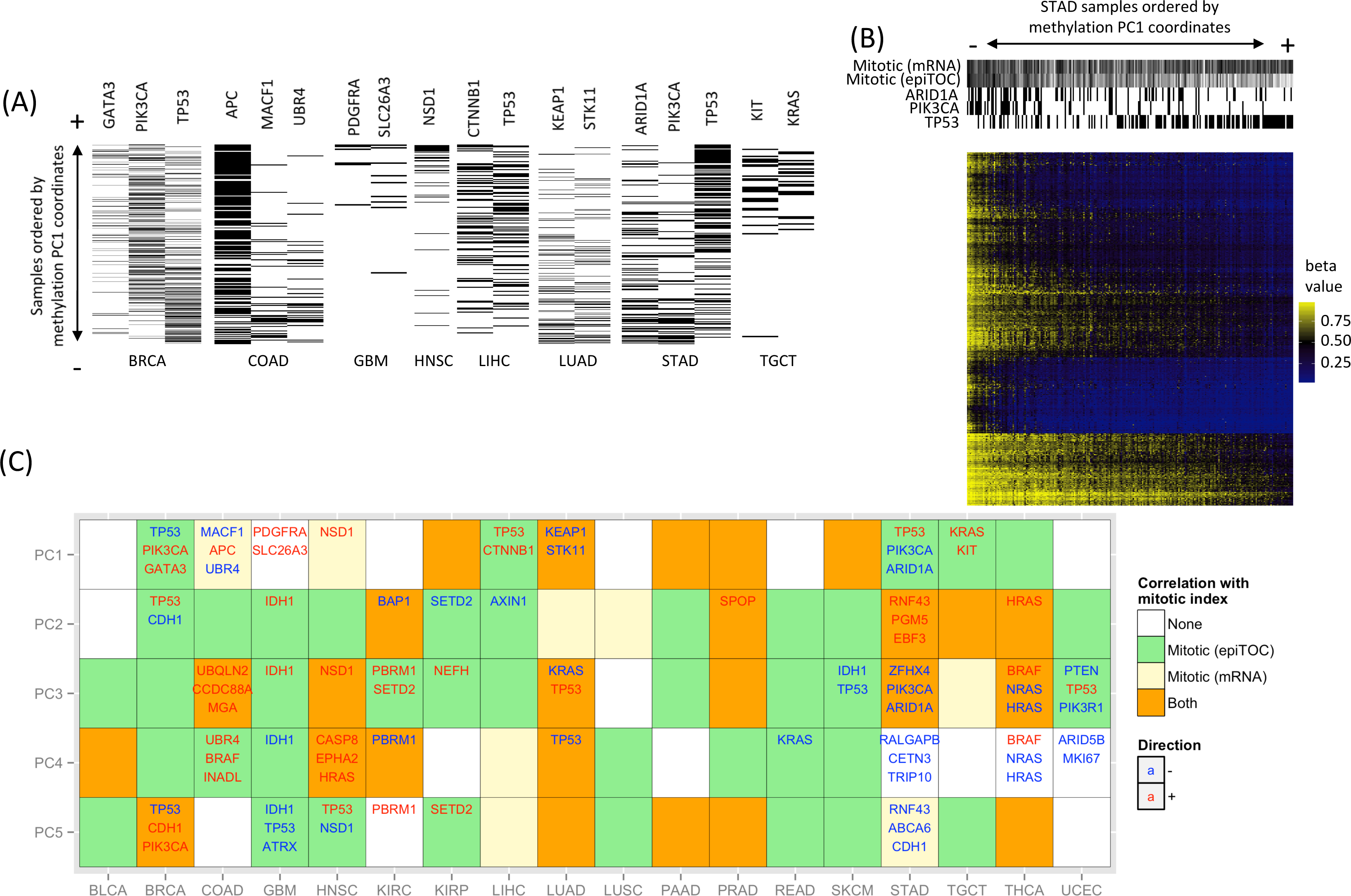
Driver gene mutations are significantly associated with DNA methylation in various cancers. (A) Examples of mutations in 15 driver genes display an uneven distribution along the first principal component (PC1) of DNAm, biased toward either the positive extreme (+) or the negative extreme (-). Tumor samples are ordered vertically by their coordinates on PC1, from small (-, *bottom*) to large (+, *top*). A black line indicates the presence of the mutated driver gene in a sample, whereas a white line indicates its absence. Note that a sample at an extreme (+/-) of a PC does not necessarily correspond to high or low methylation. (B) Example of a cancer with driver gene mutations unevenly distributed on PC1, resulting in distinct methylation patterns: *ARID1A/PIK3CA*-mutated stomach adenocarcinomas (STADs) display a methylation pattern distinct from *TP53*-mutated STADs. Shown is a heat map of methylation levels for the top 1,000 most heavily weighted probes in PC1. Each column represents a sample ordered by its PC1 coordinate, from small (-, *left*) to large (+, *right*). Each row represents a probe site. Five column sidebars are shown at the top. The bottom three indicate mutation statuses for *TP53, PIK3CA*, and *ARID1A. TP53*-mutated STADs display lower methylation levels at the selected CpG sites compared with the majority of *ARID1A*/*PIK3CA*-mutated STADs. The top two indicate cell proliferation rates estimated by the DNAm-based mitotic index (epiTOC) and the mRNA expression-based mitotic index [7]. Each index is normalized to the range between 0 (white) and 1 (black). Samples ordered by PC1 are significantly correlated with epiTOC (q=0) but not the mRNA expression-based index (q=0.22) [see results in (C)]. (C) In 15 of 18 cancer types examined, mutated driver genes were associated with one or more of the top five methylation PCs, shown as rows. The three driver genes most significantly associated with each PC are reported. Driver genes associated with a negative extreme of the PC are blue, whereas associations with the positive extreme are written in red. Background colors indicate correlations (q<0.05; Spearman correlation) with two mitotic indices: DNAm-based index (epiTOC) (light green), or mRNA-based index (light yellow), or both (orange). See Table 1 for cancer type abbreviation.

**Table 1.** Number of tumor samples, control samples, and driver genes across 18 cancer types

| Cancer type | # tumor samples | # control samples | # driver genes<sup>a</sup> |
| --- | --- | --- | --- |
| Bladder urothelial carcinoma (BLCA) | 130 | 21 | 48 |
| Breast invasive carcinoma (BRCA) | 652 | 102 | 51 |
| Colon adenocarcinoma (COAD) | 219 | 38 | 650 |
| Glioblastoma multiforme (GBM) | 144 | 2 | 16 |
| Head and neck squamous cell carcinoma (HNSC) | 306 | 60 | 34 |
| Kidney renal clear cell carcinoma (KIRC) | 245 | 160 | 14 |
| Kidney renal papillary cell carcinoma (KIRP) | 153 | 45 | 10 |
| Liver hepatocellular carcinoma (LIHC) | 202 | 50 | 11 |
| Lung adenocarcinoma (LUAD) | 407 | 32 | 84 |
| Lung squamous cell carcinoma (LUSC) | 74 | 43 | 10 |
| Pancreatic adenocarcinoma (PAAD) | 90 | 10 | 31 |
| Prostate adenocarcinoma (PRAD) | 261 | 50 | 22 |
| Rectum adenocarcinoma (READ) | 80 | 7 | 7 |
| Skin cutaneous melanoma (SKCM) | 372 | 3 | 56 |
| Stomach adenocarcinoma (STAD) | 244 | 2 | 366 |
| Testicular germ cell tumor (TGCT) | 142 | 0 | 25 |
| Thyroid carcinoma (THCA) | 446 | 56 | 5 |
| Uterine corpus endometrial carcinoma (UCEC) | 135 | 46 | 62 |

<sup>a</sup> Defined as MutSigCV-reported driver genes mutated in at least five tumor samples from our data set for a given cancer type (see MutSigCV in [17]).

A PC-associated driver gene suggests that the mutated samples at one extreme display methylation patterns distinct from the non-mutated samples at the other extreme. For instance, in stomach adenocarcinoma (STAD), *TP53*-mutated samples were distributed toward the positive extreme of PC1, whereas *ARID1A*- and *PIK3CA*-mutated samples were distributed toward the negative extreme. Thus, distinct methylation patterns associated with PC1 separated the majority of *TP53*-mutated STAD samples from *ARID1A*- and *PIK3CA*-mutated samples: *ARID1A*- and *PIK3CA*-mutated samples were highly methylated at PC1-defining probes relative to *TP53*-mutated samples (Fig 1B). Overall, we found a significant association between 159 driver genes and one or more of the top five methylation PCs in 15 of 18 cancer types (the top three driver genes associated with each PC shown in Fig 1C).

Mathematically, distinct PCs represent mutually orthogonal (uncorrelated) linear combinations of probes, or different methylation patterns (such as the pattern associated with PC1 in STAD shown in Fig 1B), with several top PCs usually capturing the majority of variance in the methylome. Thus, frequent driver gene-PC associations in almost every cancer type suggest a tight connection between driver-gene mutations and DNA methylation alterations in cancer.

Next, we investigated whether the mutation-methylation connection in cancer was limited to certain CpG subsets—namely those occurring in CGIs, shores and shelves (SSs; the 4-kb regions flanking the CGIs), or open-sea regions (i.e., outside of CGIs and SSs)—as the regulatory functions of these CpG subsets often differ [18]. For example, DNAm in promoter CGIs often causes gene silencing whereas DNAm in CGI shores is frequently altered and strongly correlated to corresponding gene expression in cancer [19]. Thus, we repeated the analysis described above for each subset of probes. We observed similar driver gene-PC associations across multiple cancer types, indicating that the mutation-methylation connection is not limited to a particular CpG subset (S1 Fig). In total, 14 of 18 cancer types harbored significant associations between driver gene mutations and top five methylation PCs at CGIs, 14 of 18 at SSs, and 15 of 18 in open sea regions. We then repeated the same analysis after stratifying probes by hypo-or hypermethylation status and found that the results did not vary appreciably (S1 Fig). Of note, in this study we identified hyper- and hypomethylated probes by comparing the methylation of tumor and control samples (q<0.05; Wilcoxon rank sum test).

### Driver gene mutations, methylation patterns, and cell proliferation are correlated

Cell proliferation was proposed to be an important factor driving DNAm changes in cancer [7]. This was demonstrated with a set of 385 CpGs in the Illumina 450K array whose average methylation levels approximated the mitotic rate in cancer, called epiTOC (epigenetic Timer Of Cancer). The method has been proposed as an alternative to the mRNA expression-based mitotic index [7]. We calculated the mitotic index for all tumor types using epiTOC or the mRNA expression-based approach. We found these two indices were correlated with many of the top 5 PCs across cancer types (q<0.05; Spearman correlation) (Fig 1C). In general, epiTOC correlated with more PCs than the mRNA expression-based index. For example, STAD showed a significant correlation between PC1 and epiTOC scores (q=0) but not the mRNA expression-based index (q=0.22) (Fig 1B). We removed epiTOC-correlated methylation sites (q<0.05; Pearson correlation), and found that many methylation PC associated-driver genes remained in many cancer types (S2 Fig), indicating that the mutation-methylation association cannot be totally explained by epiTOC scores. We also used both methods to compare predicted cell proliferation rates. To elucidate the relationship between cell proliferation rate and mutated driver genes, we associated the presence of mutations in each driver gene with high/low score in each index. We found that mutated driver genes associated with high and low cell proliferation rates estimated by the two indices were inconsistent in most cancer types (Table 2). For example, *TP53*-mutated tumors were correlated with a high cell proliferation rate in 10 cancer types and none were correlated with low cell proliferation rate using the mRNA expression-based index. In contrast, *TP53*-mutated tumors were correlated with low cell proliferation in HNSC and UCEC, using epiTOC. Thus DNAm and mRNA-based mitotic indices are inconsistent and could be measuring two different properties of these tumor cells. Despite their inconsistency, by using either index, our results indicate that methylome patterns in cancer are correlated with the presence of mutated driver genes and cell proliferation but the mutation-methylation association found in this study cannot be explained by cell proliferation alone.

**Table 2.**
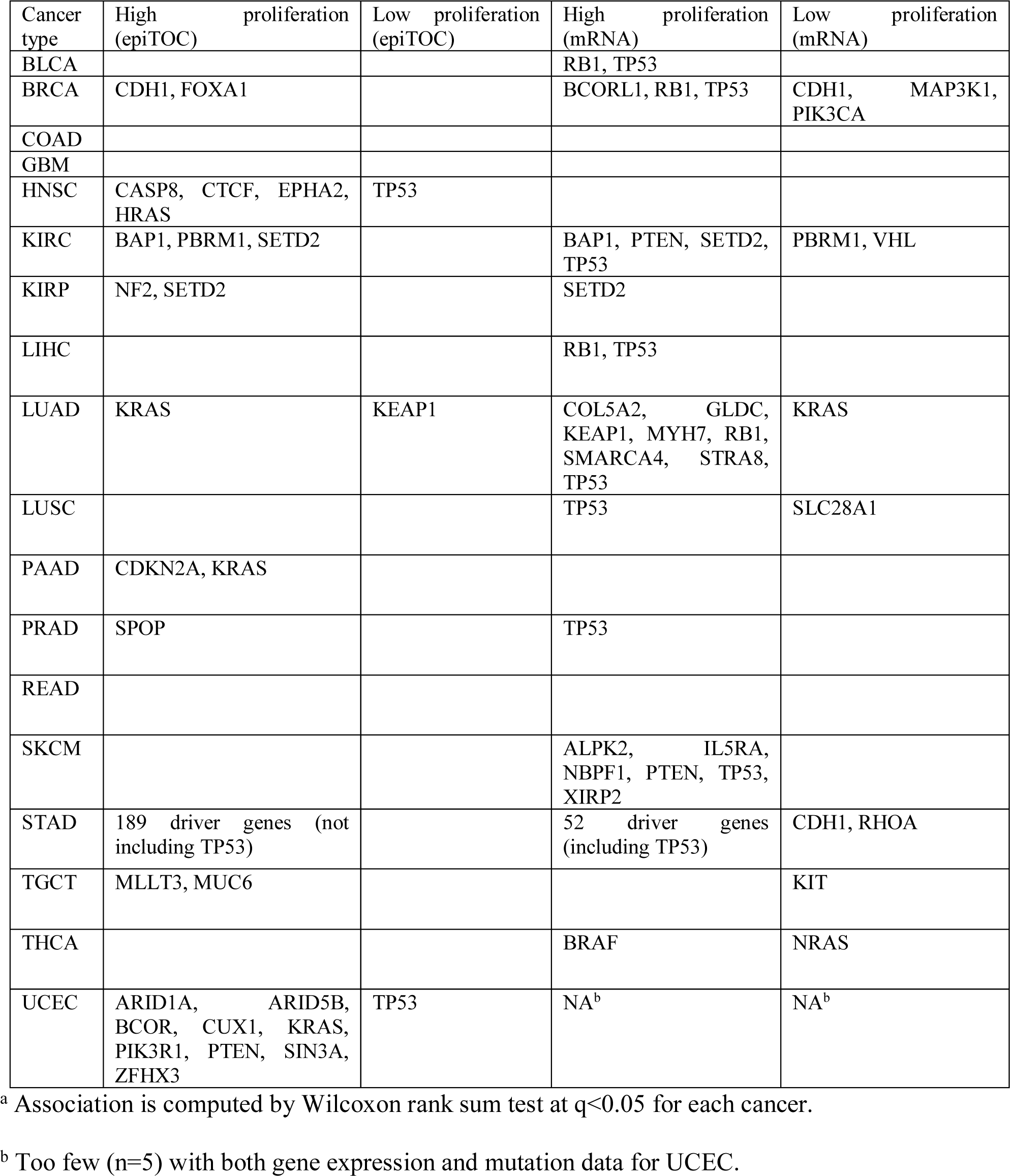
Cell proliferation rate and associated mutated driver genes

### Driver gene mutations, genome-wide CGI hypermethylation, and open sea hypomethylation in tumors

We next asked whether driver gene-associated methylation alterations corresponded to genome-wide methylation patterns characteristic of cancer: i.e., widespread CGI hypermethylation and huge hypomethylated blocks, primarily in open sea regions [1, 2, 20]. To answer this question, we calculated the HyperZ and HypoZ indices for each sample [21]. A high HyperZ index indicates aberrant hypermethylation in many CGIs for a given sample, whereas a high HypoZ index indicates extensive open sea hypomethylation. The number of mutated driver genes that were significantly associated with either high or low HyperZ and/or HypoZ indices is shown for all 18 cancer types in Table 3 (q<0.05; Wilcoxon rank sum test); it varies from 67 driver genes in COAD to 0 in PAAD, READ, and SKCM.

**Table 3.**
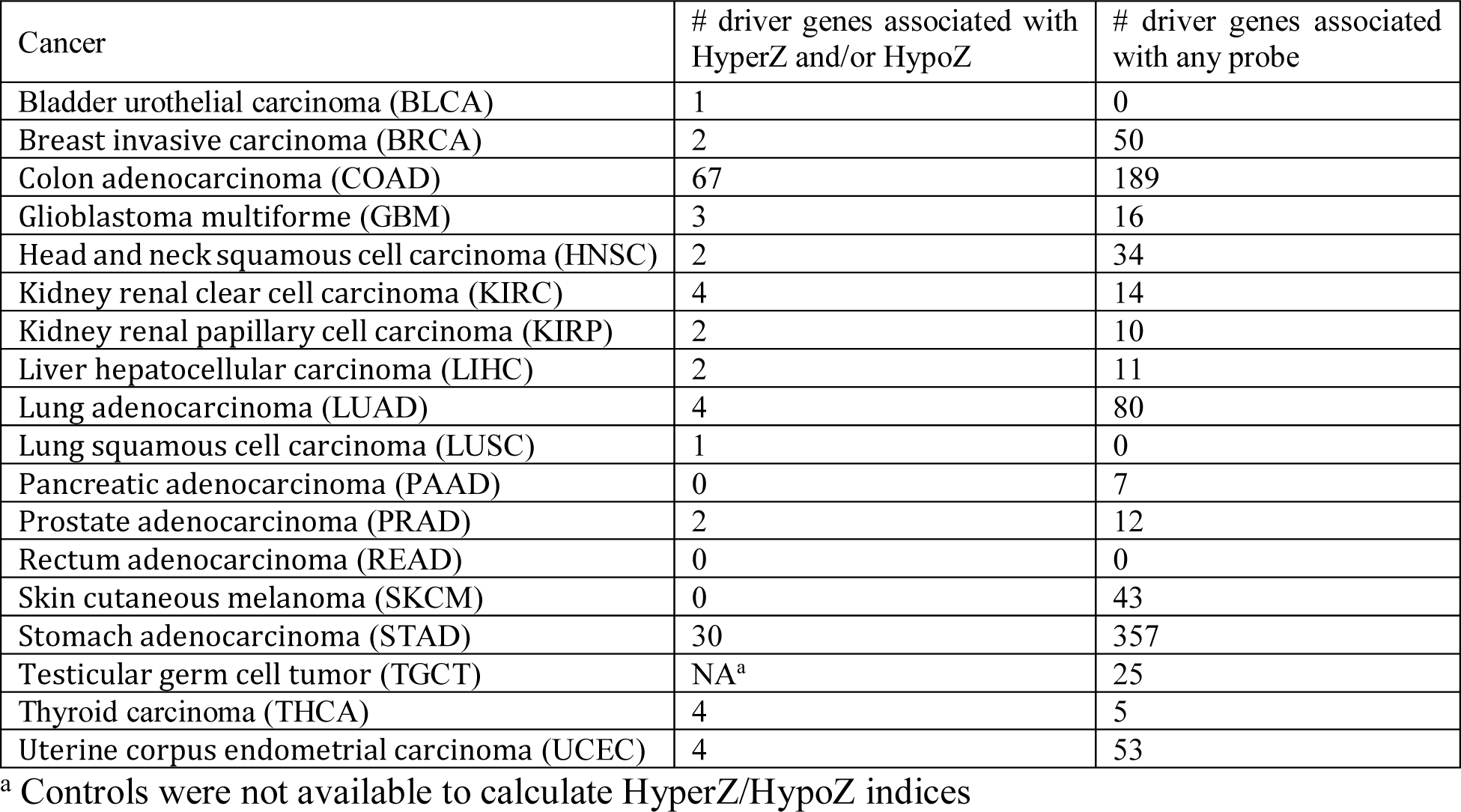
Associations between mutated driver genes and HyperZ and HypoZ indices or site-specific methylation alterations.

The complete list of mutated driver genes with significant associations to HyperZ and HypoZ is shown in Table S1. Some known players appear on this list. For example, a high HyperZ index was associated with *BRAF* in COAD and *IDH1* in GBM; both genes are linked to CIMP in cancer [13, 22]. *NSD1*, which was associated with a high HypoZ index in HNSC, has also been linked to hypomethylation in cancer [23]. The associations we detected in most cancer types underscore the relationship between driver gene mutations and genome-wide methylation alterations commonly observed in cancer, whereas only a few types lack these patterns.

### Driver gene mutations and site-specific methylation changes in tumors

Next, we investigated whether the connection between driver gene mutations and methylation alterations was methylation-site specific, in each cancer type. To do so, we calculated the associations between every driver gene and every methylation array probe for all 18 cancer types, testing whether the presence of mutations in a driver gene is associated with the high or low methylation level at a given probe site. Across almost all cancer types, many more driver genes were significantly associated with at least one probe than with the HyperZ and/or HypoZ indices, after correcting for multiple testing (*q* < 0.05; Wilcoxon rank-sum test; Table 3). In total, 737 unique driver genes were implicated. The driver gene-methylation site associations were present genome-wide. An example of the chromosomal distribution of driver gene-associated methylation probes present in kidney renal clear cell carcinoma (KIRC) is shown in Fig S3. The numerous gene-probe associations detected here suggest that driver gene-associated methylation changes likely occur at certain CpG sites, potentially resulting from a site-targeting mechanism.

The number of probes associated with each driver gene varied greatly, ranging from fewer than 10 to tens of thousands (S2 Table). For each cancer type, a few (1–5) dominant driver genes accounted for the majority of associations (Fig 2A), including known oncogenes and tumor suppressor genes such as *TP53, PTEN*, and *PIK3CA*, and known CIMP-driving genes such as *BRAF, IDH1*, and *KRAS* [15, 16, 24]. Dominant driver genes usually displayed both positive and negative associations with methylation levels in a given cancer type (Fig 2B). However, there was typically more of one type of association than the other. By definition, positive associations indicate higher methylation levels in the presence of driver gene mutations among tumor samples, whereas negative associations indicate lower methylation levels. Thus, positive associations would correspond to hypermethylation (primarily in CGIs) if control samples displayed the low methylation level at a given probe site, whereas negative associations would correspond to hypomethylation (primarily in open-sea regions) if control samples displayed the high methylation level. We found both positive and negative associations for many driver genes; however they can be largely imbalanced. For example primarily positive associations occurred for the CIMP-driving genes *BRAF* in COAD (5,166 positive vs. 103 negative associations) and *IDH1* in GBM (31,877 vs. 3,535). Other genes with predominantly positive associations included *RNF43* (4,784 vs. 173) and *MACF1* (2,706 vs. 95) in COAD; *CASP8* (21,552 vs. 4,660) in HNSC; *PBRM1* (11,831 vs. 3,906), *SETD2* (6,443 vs. 2,440), and *BAP1* (5,162 vs. 1,240) in KIRC; *SETD2* (4,300 vs. 936) in KIRP; *SPOP* (8,919 vs. 6,444) in PRAD; *PIK3CA* (58,843 vs. 2,153) in STAD; and *PTEN* (20,548 vs. 14,496) in UCEC. These genes were also associated with a high HyperZ index, suggesting that they may play a role in genome-wide CGI hypermethylation in particular cancer types (S1 Table).

**Fig 2.**
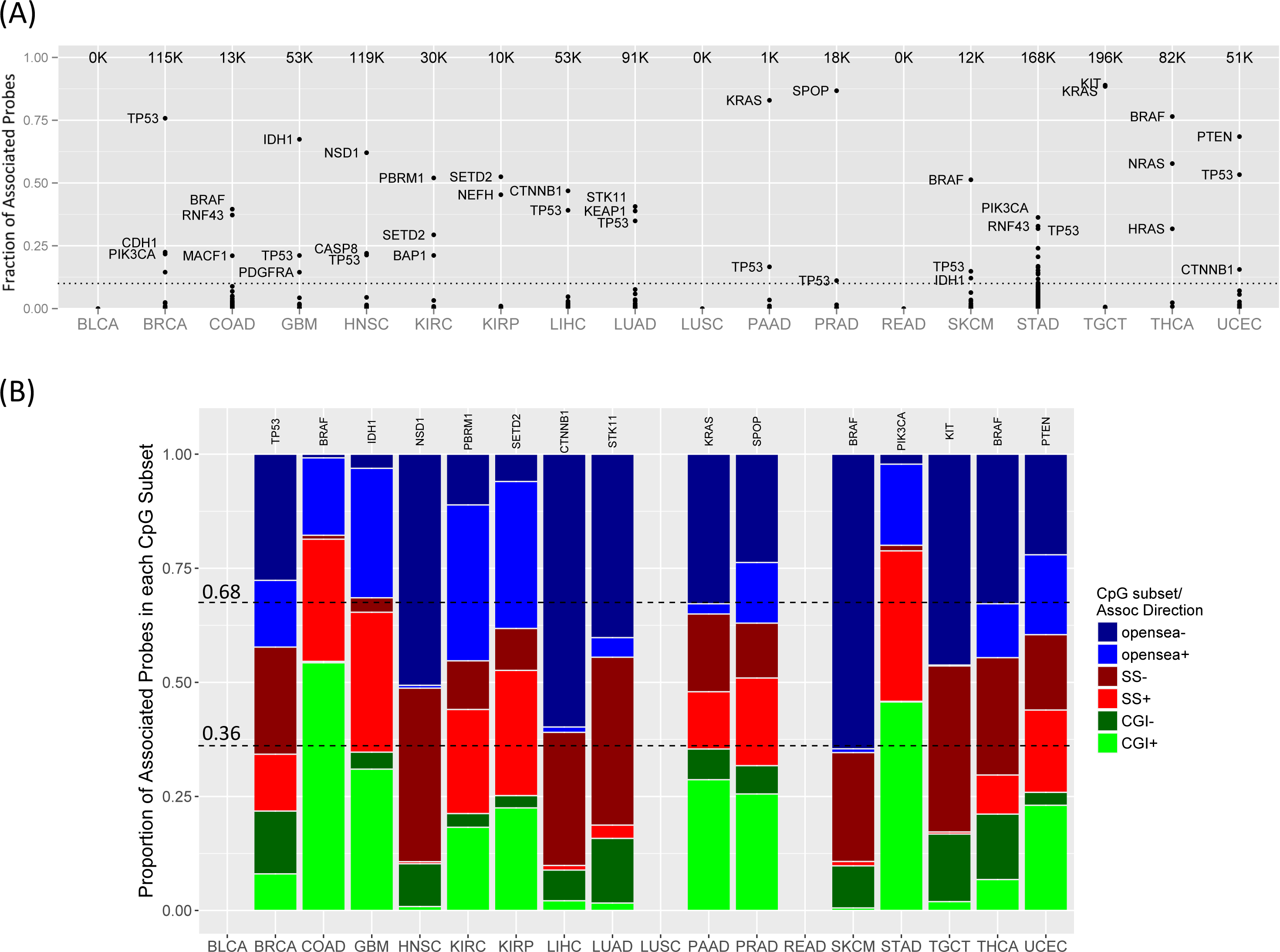
Driver gene-methylation associations and CpG subsets. (A) The total number of probes associated with any driver gene is shown for each cancer type (*top of each column*). Each point represents the fraction of corresponding probes associated with a driver gene (y-axis). Names are shown for the top three driver genes if they account for more than 10% of total probes (dotted line). (B) Driver genes with the most probe associations in each cancer type (gene names in panel). The bar plots show the proportion of associated probes in each of the three CpG subsets (CGIs; SSs; or open sea), stratified by the direction of association (+/-). Dashed lines indicate divisions expected if associations were proportionally distributed. No probes were associated with driver genes in BLCA, LUSC, and READ. See Table 1 for cancer type abbreviations.

By contrast, genes that primarily displayed negative associations were often associated with a high HypoZ index, suggesting that they may play a role in genome-wide open sea hypomethylation. Examples were *NSD1* (1,423 positive vs. 72,475 negative associations) in HNSC, *CTNNB1* (1,065 vs. 23,869) in LIHC, and *STK11* (3,269 vs. 33,840) and *KEAP1* (4,122 vs. 31,348) in LUAD. Interestingly, *RNF43* (31,905 vs. 23,267) was associated with both high HyperZ and HypoZ indices in STAD, suggesting a dual role in genome-wide CGI hypermethylation and open sea hypomethylation.

In short, a few driver genes were linked to genome-wide patterns of CGI hypermethylation and open sea hypomethylation in particular cancer types, whereas many more driver genes were linked to only a few probe sites that exhibited hyper- and hypomethylation in cancer (S2 Table).

### Consistency of mutated driver gene-associated methylation changes across cancer types

Next, we asked whether mutated driver genes consistently displayed predominantly positive/negative associations across multiple cancer types. We investigated the proportion of positive and negative probe associations for 17 driver genes across 18 cancer types (Fig 3A). Here, we only considered driver genes linked to extensive methylation changes (more than 1,000 probe associations per driver gene, for at least two cancer types). When compared to control samples, positive associations often equated to hypermethylation (primarily in CGIs) in response to mutations. Likewise, the negative associations often equated to hypomethylation (primarily in open sea regions) (Fig 3B). *TP53* displayed predominantly negative associations in 9 of 18 cancer types, and no predominantly positive associations were observed in connection to this gene in any cancer type, suggesting a tight connection between mutations in this gene and open sea hypomethylation across multiple cancer types. Interestingly, those negatively associated probes were shared across cancer types (from 62,951 probes shared across 2 types to 15 probes shared across 7 types) (S4 Fig). In addition to *TP53, APC* and *CTNNB1* also displayed predominantly negative associations in two different cancer types each.

**Fig 3.**
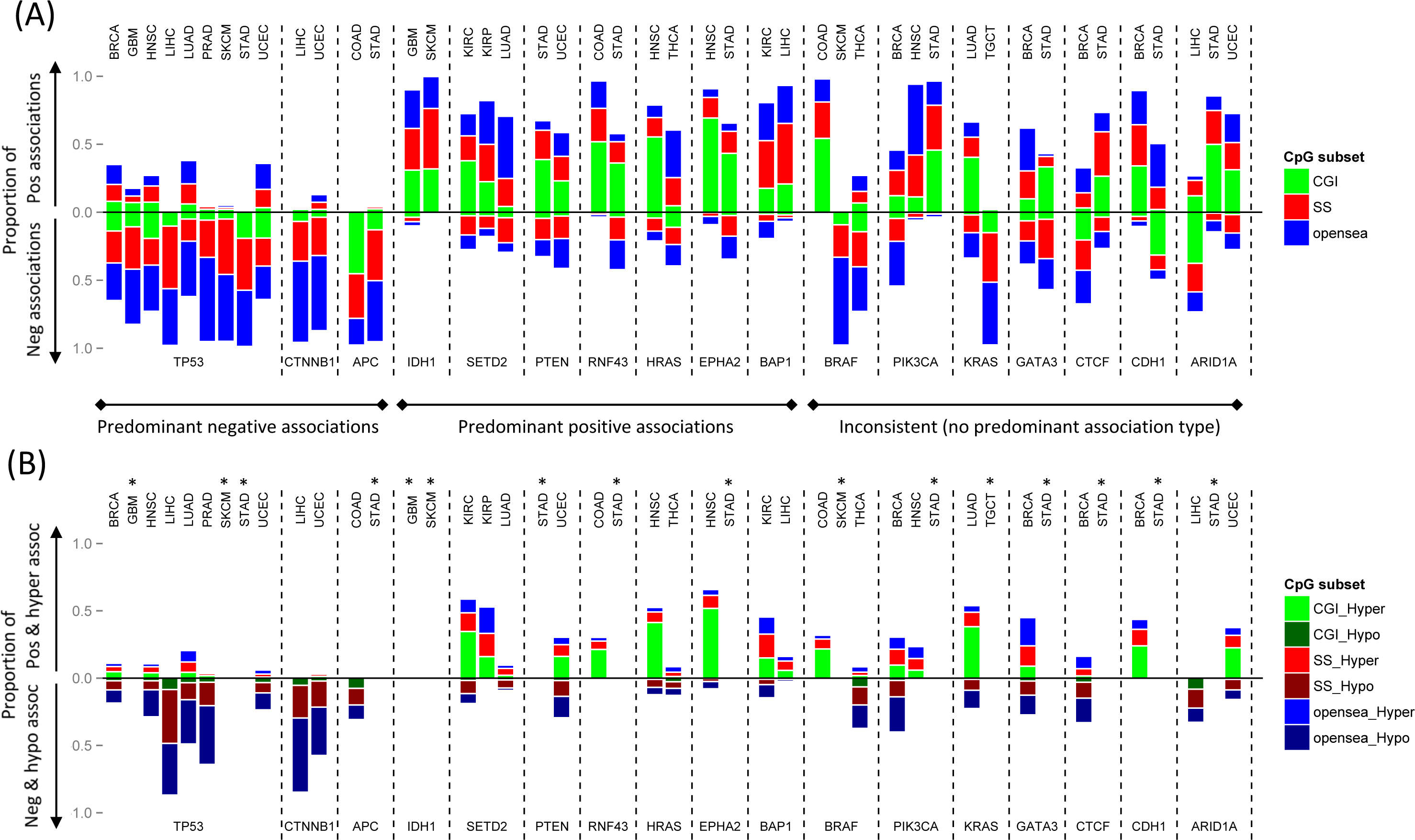
Proportions of positive and negative associations with methylation for 17 recurrently mutated driver genes. (A) Barplots show the proportion of methylation probes for each driver gene (*labels at bottom*) and cancer type (*labels at top*) displaying positive and negative associations. Positive associations are plotted above the horizontal line, negative associations are below the horizontal line. Associations are further stratified by CpG subset: CGI (CpG islands), SS (shores and shelves of CGIs; 4kb regions flanking CGIs), and open sea (regions outside CGIs and SSs). Driver genes can be classified into 3 groups based on directionality of their predominant associations (*negative, positive, inconsistent*). Genes in figure were associated with more than 1,000 probes, in at least two cancer types. (B) Plotted as in (A), using: (1) positively associated and hypermethylated probes and (2) negatively associated and hypomethylated probes. ^*^Hyper-or hypomethylated probes were not identified for GBM, STAD, SKCM, and TGCT due to a lack of control samples. See Table 1 for cancer type abbreviations.

By contrast, *IDH1* strongly favored positive associations in two cancer types, GBM and SKCM, consistent with reports that mutated *IDH1* downregulates TET-dependent demethylation, resulting in aberrant CGI hypermethylation [25]. *SETD2, PTEN, RNF43, HRAS, EPHA2*, and *BAP1* were also linked to primarily positive associations in more than one cancer type, suggesting that they may play a general role in CGI hypermethylation.

*BRAF*, which mediates CIMP in colorectal cancer, displayed a high proportion of positive associations (0.98) in COAD, but low proportions in SKCM (0.02) and THCA (0.27). Its negative associations in THCA corresponded to hypomethylation across CGIs, SSs, and open sea regions (Fig 3B and S2 Table). This dramatic difference indicates that driver genes may be associated with methylation patterns in a cancer type-specific manner. Such cancer type-specific associations were also seen for *PIK3CA, KRAS, GATA3, CTCF, CDH1*, and *ARIDA1*.

### Identification of molecular subtypes in thyroid carcinoma and uterine corpus endometrial carcinoma based on driver gene-associated methylation patterns

In previous studies, researchers have identified cancer subtypes in COAD and GBM by matching mutational profiles to methylation patterns [13, 22]; here, we asked if the site-specific mutation-methylation associations would separate THCA and UCEC tumors into subtypes. We focused on the methylation patterns associated with the top three dominant driver genes which account for the most probe associations in each cancer type (as shown in Fig 2A). In THCA, the top three genes (*NRAS, HRAS*, or *BRAF*) were mutated in a mutually exclusive fashion. However, in UCEC, mutations in *TP53* were nearly mutually exclusive with *PTEN* and *CTNNB1* mutations, which co-occurred in many tumor samples.

We performed hierarchical clustering on the union of the 500 methylation probes most significantly associated with mutations in each of the top three genes (Fig 4). In THCA, two methylation subtypes emerged corresponding to *NRAS*- and *HRAS*-mutated tumors vs. *BRAF*-mutated tumors (Fig 4A). Furthermore, we found tumors lacking any of the specified mutations co-clustered within these patterns. The selected probes primarily fell in non-CGI positions (i.e. SS and open sea regions); *BRAF* mutants displayed hypomethylation in open sea and some SS regions, whereas *NRAS* and *HRAS* mutants displayed methylation levels similar to controls in open sea and SSs regions, with little hypermethylation. Two methylation subtypes were also identified in UCEC, this time corresponding to *TP53* vs. *PTEN* mutations (Fig 4B). *PTEN-* mutated samples generally exhibited CGI hypermethylation, whereas *TP53*-mutated samples generally exhibited normal methylation levels, with some hypomethylation in open sea regions. Most UCEC samples with mutations in both *PTEN* and *CTNNB1* displayed greater levels of open sea hypomethylation than samples with *PTEN* mutations alone, which has not been previously reported, illustrating the connectivity between mutational profiles and DNA methylation in cancer.

**Fig 4.**
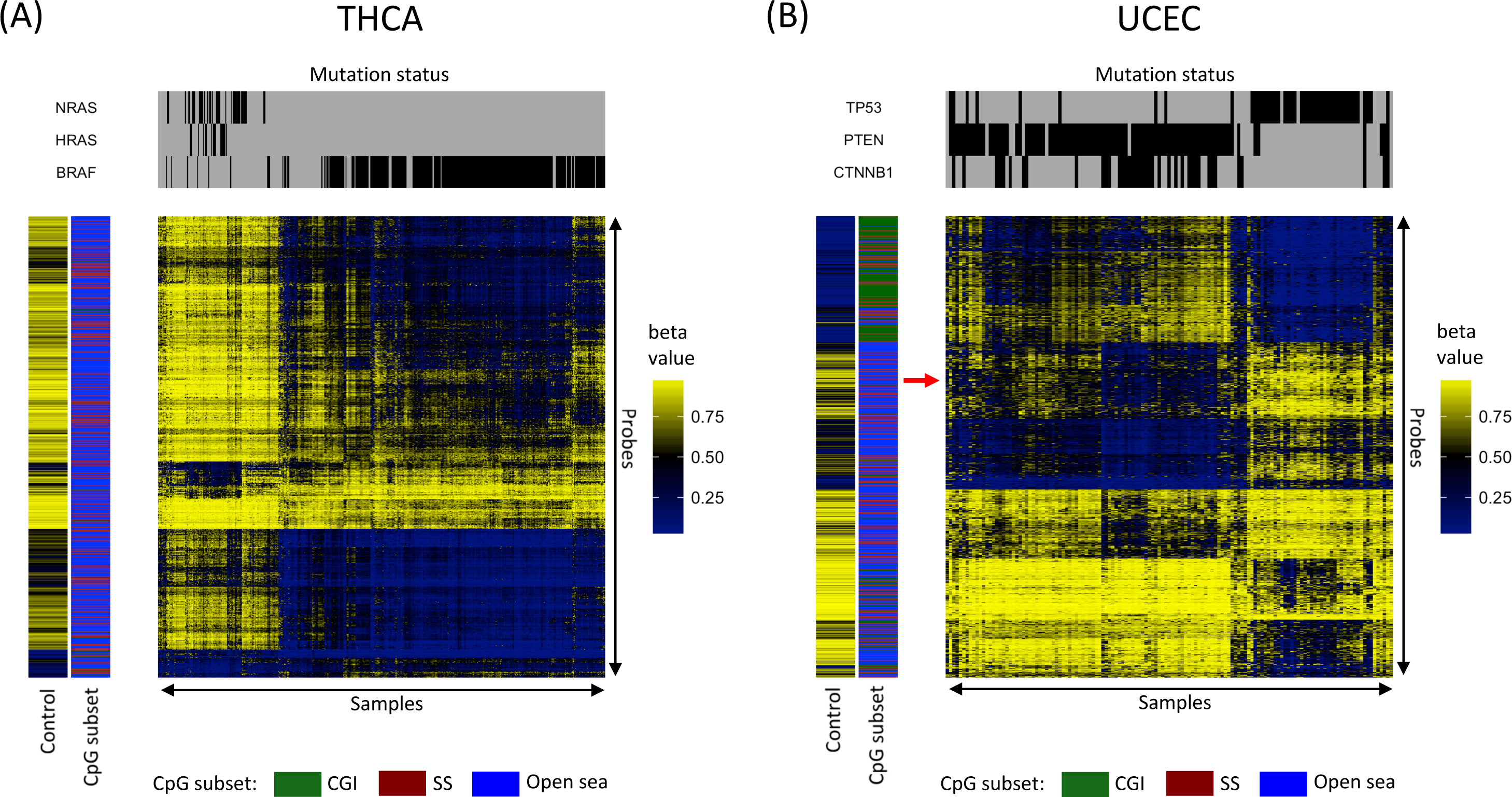
Driver gene-associated methylation patterns can be used to identify molecular subtypes. Heat maps for (A) thyroid carcinoma (THCA) and (B) uterine corpus endometrial carcinoma (UCEC) depict hierarchical clustering of methylation values of the union set of the 500 probes most significantly associated with each of the three dominant driver genes in each cancer type. Each column represents a sample, and each row represents a probe. Mutation status shown in upper panel. Left panel sidebars indicate CpG subset and average methylation level across control samples. The arrow in (B) indicates methylation probes displaying more hypomethylation in samples where *PTEN* and *CTNNB1* mutations co-occurred than samples with *PTEN* mutations alone.

### Correlations between driver gene-associated methylation probe sites and corresponding gene expression in two thyroid carcinoma subtypes

Finally, we investigated whether driver gene-associated methylation patterns shape gene regulatory networks. To investigate the three-way association between mutation, methylation, and gene expression, we used THCA as our primary example; the mutually exclusive mutation profile present in this type of cancer minimized the complexity of the associated methylation patterns, facilitating the study of gene expression. We looked for genes whose aberrant expression levels were correlated with aberrant methylation levels (each relative to controls). We focused on CpG sites in promoter regions and gene bodies in *NRAS* and *HRAS* mutants (the *NRAS-HRAS* group) vs. *BRAF* mutants (the *BRAF* group). These genes were subsequently sorted into four different categories of expression change: (1) upregulation only in the *BRAF* group, (2) upregulation only in the *NRAS*-*HRAS* group, (3) downregulation only in the *BRAF* group, and (4) downregulation only in the *NRAS*-*HRAS* group. For each category, we reported the affected genes, sorted by median difference in expression between mutants and controls (S3 Table), and significantly enriched pathways (S4 Table).

In all, 1,565 genes were upregulated specifically in the *BRAF* group (S3 Table). Gene set analysis showed that these genes were enriched in 97 canonical pathways primarily pertinent to cell-cell communication/extracellular matrix gene sets and signaling pathways (q<0.05; hypergeometric test; S4 Table). Some upregulated genes were involved in many of the 97 pathways; for example, the 10 genes implicated in the most pathways were: *GRB2* (present in 34 out of 97 pathways), *RAC1* (*n*=26), *STAT1* (*n*=25), *LYN* (*n*=24), *VAV1* (*n*=24), *JAK1* (*n*=22), *PTPN6* (*n*=22), *ITGB1* (*n*=20), *STAT3* (*n*=18), and *STAT5* (*n*=18). We noticed that genes in JAK and STAT gene families (e.g. *STAT1, JAK1, STAT3*, and *STAT5*) were differentially regulated between two THCA subtypes and implicated in many signaling cascades, including the KEGG JAK-STAT signaling pathway (ranked 12^th^ out of the 97 pathways; *q*=4.7E-5). In addition, *JAK3* (*n*=11) and *STAT4* (*n*=4), were also upregulated and present in multiple pathways. Our results show that this differential regulation of the JAK and STAT families may be shaped by differences in DNA methylation (Fig 5A). Specifically, *STAT1* differential expression is negatively correlated with methylation levels at the SS probe of the promoter CGI, whereas *JAK3* differential expression is positively correlated with methylation levels at the 3´ gene body CGI and its north shore (Figs 5B and 5C). In addition, 9 of the top 15 differentially methylated genes were involved in metastasis: *KLK6, KLK7, KLK11* [27], *CLDN10* [28], *B3GNT3* [29], *RASGRF1* [30], *ST6GALNAC5* [31], *TACSTD2* (also known as *TROP2*) [32], and *CEACAM6* [33] (S3 Table).

**Fig 5.**
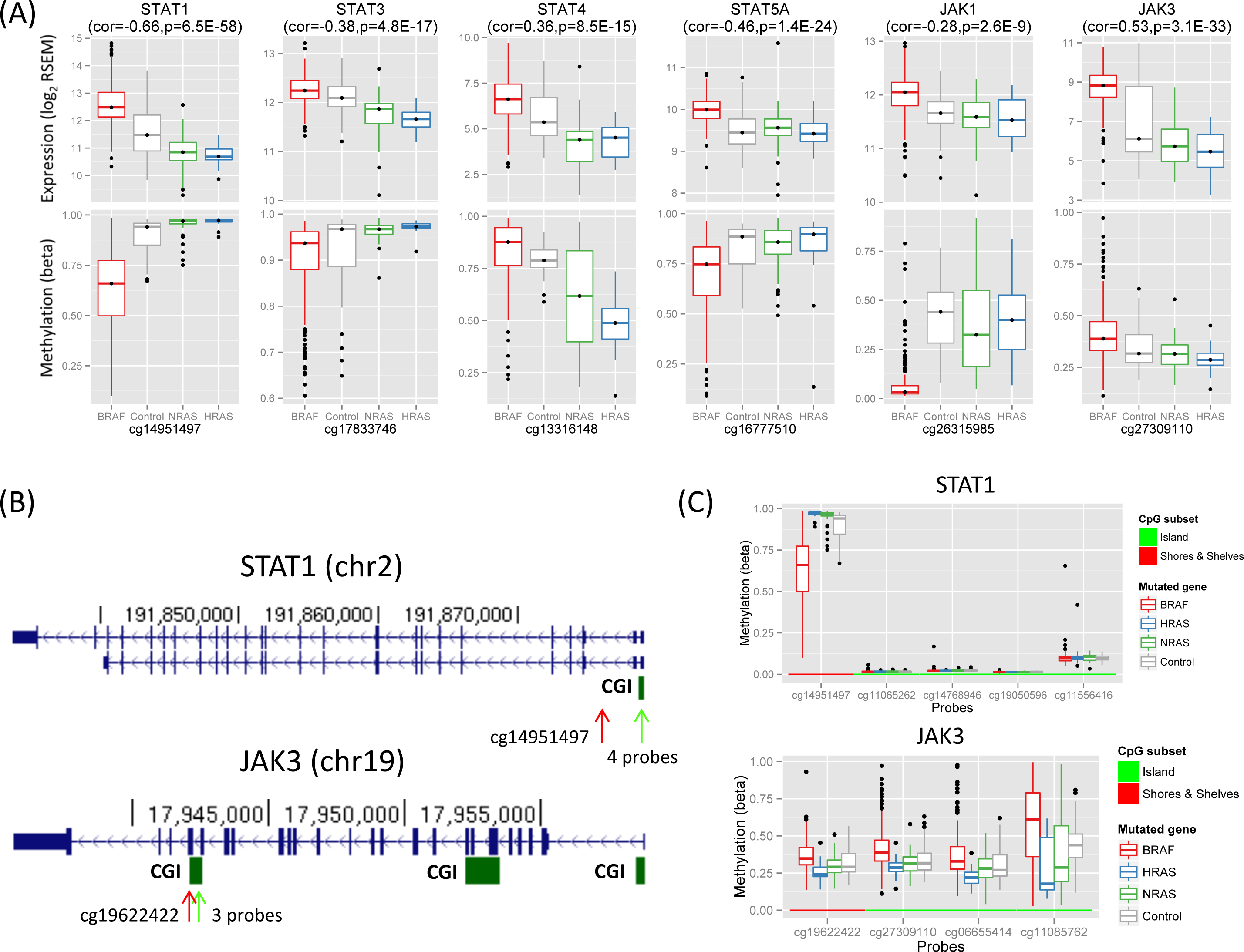
Differential expression of JAK and STAT family genes correlated with differential methylation in thyroid cancer. (A) Shown are box plots for gene expression levels (*top plots*) and methylation levels (*bottom plots*) for *STAT1, STAT3, STAT4, STAT5A, JAK1*, and *JAK3*, grouped by *BRAF*-mutated (*red*), *HRAS*-mutated (*blue*), *NRAS*-mutated (*green*), and control (*grey*) samples. Gene names and Spearman rho (with p-value) between gene expression and methylation among tumor samples are shown on top. Probe names (where methylation levels were measured) are shown at bottom. (B) Shown are snapshots from the UCSC genome browser for *STAT1* and *JAK3*, with CpG islands (CGIs) indicated below (*green arrow*: CGIs; *red arrow*: probes flanking the CGIs). Methylation levels of the indicated regions are shown in panel (C). (C) Box plots show methylation levels (y-axis) at probes in *STAT1* and *JAK3* for *BRAF*-mutated, *HRAS*-mutated, *NRAS*-mutated, and control samples. The shown probes fall in the south shores and shelves [or SSs, indicated by red arrows in (B)] of the *STAT1* promoter CGI and the 3´ CGI of *JAK3* [indicated by green arrows in (B)].

By contrast, 1,043 differentially methylated genes were downregulated specifically in the *BRAF* group. Gene set analysis showed that these genes were enriched in five canonical pathways pertinent to amino acid catabolism, triglyceride biosynthesis, glycerophospholipid metabolism, and nucleic acid metabolism (S4 Table).

The *NRAS-HRAS* group displayed 278 differentially methylated, upregulated genes. These genes were enriched in three canonical pathways, relevant to the neuronal system, potassium channels, and melanogenesis (S4 Table). We did not find a significant portion of differentially methylated genes implicated in tumor progression among the top differentially expressed genes (defined by median difference in expression between mutants and controls). However when we considered differentially methylated genes within the top 17 that were highly transcribed, 6 out of 17 were implicated in tumorigenesis: G protein alpha subunit (*GNAS*) [34], pyruvate dehydrogenase kinase 4 (*PDK4*) [35], NADPH reductase (*NQO1*) [36], and three NADPH oxidases that produce H_2_O_2_ for thyroid hormone synthesis: *DUOX2, DUOXA2*, and *DUOX1* [37] (S3 Table).

By contrast, 447 differentially methylated genes were downregulated specifically in the *NRAS-HRAS* group. These genes were enriched in 166 canonical pathways; interestingly, 47 genes overlapped the 97 pathways enriched in the *BRAF* upregulated group, including the JAK-STAT signaling pathway (ranked 13^th^, *q* = 3.5E-5). Specifically, *STAT1, STAT3, STAT4*, and *JAK3* were among the genes upregulated in the *BRAF* group but downregulated in the *NRAS-HRAS* group (Fig 5A and S3 Table). This result demonstrates that methylation changes are indeed associated with differential gene expression between *BRAF*-mutated and *NRAS*- and *HRAS*-mutated samples in THCA.

## Discussion

In this study, we demonstrated that driver gene mutations are tightly tied to the DNAm landscape in multiple types of cancer. Furthermore, we showed that mutated driver genes are associated with DNAm alterations in a reproducible, site-specific manner. In each cancer type, a few driver genes dominate the site-specific associations, and some potentially contribute to CGI hypermethylation and extensive hypomethylation, i.e., the hallmarks of cancer. However, we caution that these findings do not equate to causality, but point to the highly interconnected nature of the genome and epigenome.

Our findings are consistent with previous research on methylation in cancer; however, they also contribute novel insights. For example, several driver genes that display primarily positive or negative methylation probe associations in this study have been linked to CGI hypermethylation or open sea hypomethylation, respectively. Driver genes associated with CGI hypermethylation in both this study and past studies include *BRAF* in COAD [22], *IDH1* in GBM [13], *SETD2* in KIRC [38], *PIK3CA* in STAD [3], and *PTEN* in UCEC [39], whereas genes associated with hypomethylation include *TP53* in LIHC [12] and *NSD1* in HNSC [23]. In addition to these examples, we identified novel driver genes that may contribute to CGI hypermethylation, such as *CASP8* in HNSC and *SPOP* in PRAD, or to open sea hypomethylation, such as *CTNNB1* in LIHC and *BRAF* in THCA. By illuminating the driver genes associated with widespread DNAm alterations, as well as driver genes associated with more limited DNAm alterations, our comprehensive analysis provides a detailed mutation-methylation map for many types of cancer.

Several mutated driver genes displayed consistent and widespread positive or negative associations across cancers corresponding to extensive DNAm alterations, whereas others showed effects that varied by cancer type. This discrepancy may be attributable to different underlying mechanisms. For example, mutations in *IDH1* and *SETD2* directly affect the epigenetic landscape by inhibiting TET-dependent demethylation and disturbing DNA methyltransferase targeting, respectively [14, 25, 38]. Both mechanisms cause DNA hypermethylation, in line with the corresponding primarily positive associations observed in this study. By contrast, several documented mechanisms link *TP53* loss to global hypomethylation, consistent with *TP53* mutations and hypomethylation in this study [11, 12, 40, 41]. Moreover, probes negatively associated with *TP53* were shared across cancer types, suggesting that similar CpG sites may be targeted in *TP53*-mutated tumors independent of tissue types. In contrast to patterns of consistency across cancer types, *BRAF* mutations displayed inconsistent methylation patterns between types. For example, in COAD, BRAF V600E recruits the DNA methyltransferase to CGIs targets by stimulating the MEK/ERK signaling pathway and upregulating the transcription repressor MAFG [15]. This is in line with *BRAF*-associated, widespread CGI hypermethylation in COAD observed in this study. However, in THCA, *BRAF*-mutated samples (260/266 of which harbored the V600E mutation) largely displayed hypomethylation. Although no mechanistic explanation is yet available, it could be either that the mutation does not upregulate MAFG in THCA or that MAFG is upregulated in both types resulting CGI hypermethylation at a few binding sites but some other factor is responsible for the extensive CGI hypermethylation in COAD.

It is unclear what links site-specific methylation alterations to driver gene mutations. A model of DNAm targeting proposed by Struhl [42] provides one potential mechanistic explanation. In this model, DNA methylation alterations are positioned and maintained by transcriptional circuitry that is aberrantly regulated as the result of driver gene mutations. This model was supported by the recent finding that BRAF V600E and KRAS G13D mutations in COAD upregulate the transcription factors MAFG and ZNF304, respectively, resulting in targeted promoter CGI hypermethylation near TF binding sites [15, 16]. However, in COAD, the *BRAF*-associated widespread CGI hypermethylation is not restricted to the published TF binding sites, making it unclear whether this model can be generalized to explain all driver gene-probe associations in COAD. On the other hand, several biological processes that can alter DNAm at specific sites have been documented recently. For example, cellular oxidative stress can produce hypermethylation in promoters of low-expression genes [43]. Hypoxia can reduce TET activity leading to hypermethylation at corresponding sites [44]. Cell proliferation accumulates aberrant DNAm in promoters of polycomb group target CpGs (pcgt) [7]. In this study, we showed that driver gene mutations correlate with the pcgt-derived mitotic index in many cancer types. The same connection between driver gene mutations and DNAm-altered sites may be established via other DNAm-altering processes as well.

Because mutation-methylation patterns are likely to reflect important oncogenic characteristics, using these patterns to separate tumors into molecular subtypes could potentially aid treatment selection. In a proof of concept portion of this study, we successfully identified molecular subtypes in THCA and UCEC based on the dominant driver gene-associated methylation patterns. These subtypes agreed with previous reports of subtypes defined by gene expression analyses [45, 46]. Though we only attempted to identify subtypes in two cancer types, these results indicate that our mutation-methylation-based approach could be useful for identifying molecular subtypes in other cancer types as well.

The mutually exclusive mutations in *NRAS, HRAS*, and *BRAF* in THCA tumors have been interpreted to mean that these mutations have interchangeable effects on activating MAPK signaling, the main cancer-driving event in papillary thyroid carcinomas [46]. Our analysis, however, highlights substantial differences in DNAm between *BRAF*-mutated vs. *NRAS*- and *HRAS*-mutated THCA tumors; moreover, the differences in DNAm appear to profoundly shape gene expression profiles, which may contribute to thyroid tumorigenesis. In this study, DNAm alterations in *BRAF*-mutated tumors were correlated with the activation of JAK-STAT signaling and metastasis, whereas DNAm alterations in *NRAS*- and *HRAS*-mutated tumors were linked to potential DNA damage by H_2_O_2_ overproduction [37], activation of G-protein signaling by *GNAS* overexpression [34, 47], and activation of mTOR signaling by *PDK4* overexpression [46]. These differences in the molecular processes linked to different driver gene mutations may contribute to distinct pathways of tumorigenesis, yielding different prognoses and clinical phenotypes.

This work has several limitations. First, all samples with mutations in the same gene were classified together. For example, although the majority of *BRAF*-mutated samples carried the V600E mutation (25 out of 34 *BRAF*-mutated tumors with BRAF V600E in COAD, 167/195 in SKCM, and 260/266 in THCA), this group also included a few non-V600E mutations. We took this approach because MutSigCV only reports driver genes but not mutations. However, different mutations in the same gene may be linked to different methylation patterns, introducing noise into our analysis and lowering our statistical power to detect mutation-methylation associations. Second, we examined the individual associations between driver genes and methylation sites. However, the combinatorial effect of driver gene mutations on methylation could exist, since several driver gene mutations can typically co-occur in a given tumor. Third, we focused only on MutSigCV-reported driver genes and were limited to the information present in TCGA data. Although MutSigCV is one of the most reliable driver gene detection tools available, limitations associated with the detection algorithm—paired with the limitations imposed by the number of tumor samples available in TCGA—may have led us to miss methylation-altering mutations that occurred in unknown driver genes. Finally, tumor purity is a potential confounding factor in analyses of cancer data. Although we were not able to exclude the possibility of confounding, the mutation-methylation associations reported here were seen in cancer types where most TCGA samples (>80%) were predicted with high purity (>70%), including GBM, KIRP, THCA, and UCEC [48]. Therefore, it seems unlikely that confounding by tumor purity level was extensive.

## Conclusions

This pan-cancer analysis provides the strongest evidence to date for a widespread connection between genomic and epigenomic alterations in cancer. The mutation-methylation relationships described here could potentially be used to identify tumor subtypes, thus aiding in prognosis and treatment decisions. In addition, in the future, further analysis of methylation and expression data may identify driver gene mutation-induced methylation alterations that dysregulate genes/pathways that promote tumor growth. Importantly, such dysregulation could potentially be corrected by treating patients with agents that influence the DNA methylation landscape. Demethylating agents such as 5-aza-2´-deoxycytidine, for example, have been used to reactivate epigenetically silenced tumor suppressor genes and also to decrease overexpression of oncogenes [49, 50]. By contrast, the methyl donor S-adenosylmethionine has been shown to downregulate the oncogenes *c-MYC* and *HRAS*, inhibiting cancer cell growth [51]. In summary, in light of the connection between driver gene mutations and DNA methylation shown here, it will be important to further study how coordinated genomic and epigenomic alterations result in the hallmarks of cancer. A better understanding of the molecular mechanisms underlying cancer may help us identify factors that accelerate tumor onset, predict biomarkers for early diagnosis, and assess new therapeutic targets.

## Materials and Methods

We analyzed samples that had both somatic mutation data and DNA methylation data available. These samples represented 18 cancer types: bladder urothelial carcinoma (BLCA), breast invasive carcinoma (BRCA), colon adenocarcinoma (COAD), glioblastoma multiforme (GBM), head and neck squamous cell carcinoma (HNSC), kidney renal clear cell carcinoma (KIRC), kidney renal papillary cell carcinoma (KIRP), liver hepatocellular carcinoma (LIHC), lung adenocarcinoma (LUAD), lung squamous cell carcinoma (LUSC), pancreatic adenocarcinoma (PAAD), prostate adenocarcinoma (PRAD), rectum adenocarcinoma (READ), skin cutaneous melanoma (SKCM), stomach adenocarcinoma (STAD), testicular germ cell tumors (TGCT), thyroid carcinoma (THCA), and uterine corpus endometrial carcinoma (UCEC). The number of tumors and controls (normal tissue samples adjacent to tumors or healthy donors) for each cancer type is listed in Table 1.

### Data preprocessing

Exome-sequenced level 2 somatic mutation data were downloaded from TCGA’s data portal (https://tcga-data.nci.nih.gov/tcga/) on February 1, 2015.

TCGA level 3 DNA methylation array-based data (Illumina Infinium HumanMethylation450 BeadChip array) were downloaded from the UCSC Cancer Genomics Browser (https://genome-cancer.ucsc.edu) on October 26, 2015. DNA methylation levels were measured with β values. We normalized β values for type I and II probes using the β mixture quantile (BMIQ) method [52]. The following types of probes were removed from the analysis: (1) probes on the X and Y chromosomes, (2) cross-reactive probes [53], (3) probes near single nucleotide polymorphisms (SNPs), and (4) probes with missing rates ≥90% across all samples for a given cancer type. A final set of 314,421 probes was analyzed for each cancer type.

Finally, TCGA level-3 gene expression data measured by log transformed (base 2) RSEM-normalized RNA-Seq (Illumina HiSeq) counts were downloaded from the UCSC Cancer Genomics Browser (https://genome-cancer.ucsc.edu) on November 4, 2015.

### Driver genes

We defined driver genes as those reported by MutSigCV2 [17] for each of the 18 cancer types; these data were downloaded from the Broad Institute of MIT & Harvard (http://firebrowse.org). Specifically, we analyzed genes that had reported q-values < 0.05 and that were mutated in at least five samples from each cancer type. The number of driver genes for the 18 cancer types is summarized in Table 1.

### Hyper- and hypomethylated probes

For each probe, we compared the distribution of methylation levels among tumor samples with that among control samples using one-sided Wilcoxon rank sum tests, one for each direction, stratified by cancer type. Each significant probe (*q* < 0.05) was classified as either hypermethylated (methylation levels in tumor samples were greater than in control samples) or hypomethylated (methylation levels in tumor samples were less than in control samples).

### Principal component analysis

PCA was performed using the R package ‘pcaMethods,’ and missing values were imputed by probabilistic PCA. The top five PCs were computed within each cancer type for all probes and subsets of probes, including probes in CGIs, the SSs around CGIs (the 4-kb regions flanking CGIs), and open sea regions (CpGs outside CGIs and SSs). The probe sets (CGIs, SSs, and open sea regions) were further stratified by hyper- and hypomethylation status. CGIs, SSs, and open sea regions were defined in the Illumina 450K array annotation file.

### HyperZ and HypoZ indices

HyperZ and HypoZ indices were computed for each tumor sample within a cancer type. The HyperZ and HypoZ indices were introduced by Yang et al. [21] to measure the level of overall CGI hypermethylation and open sea hypomethylation, respectively.

### Association between driver gene mutations and DNA methylation

We analyzed somatic mutations at the gene level. A driver gene was classified as either mutated (any mutations) or not mutated (no mutations) for each tumor sample. Associations between driver gene mutations and methylation were tested using the two-sided Wilcoxon rank sum test. To evaluate driver gene-PC associations, the test was performed for every driver gene and each of the top five methylation PCs; samples were ranked based on their coordinates on the PC, and the mutated cohort was compared with the non-mutated cohort. To evaluate site-specific associations, the test was performed for every possible driver gene-probe pair. Here, samples were ranked based on their β values at the probe, and the mutated cohort was compared with the non-mutated cohort. Finally, we performed the same association test for every driver gene and the HyperZ and HypoZ indices, to identify driver genes potentially associated with genome-wide CGI hypermethylation and open sea hypomethylation. In each case, q-values were computed by correcting for all tests performed for a given cancer type [54]. Associations were considered significant at *q* < 0.05. For associations between every gene-probe pair, the empirical false discovery rate was also estimated by permuting the mutation status for every driver gene. The results showed that the empirical false discovery rate was controlled (< 0.05) at the theoretical cutoff (*q* < 0.05) for each cancer type (S5 Fig).

### Correlations between methylation PCs, mutated driver genes, and mitotic indices

A DNAm-based mitotic index called epiTOC (epigenetic Timer Of Cancer) was proposed by Yang et al. to estimate tumor cell proliferation rates. Performance was validated by correlating with a mRNA expression-based mitotic index [7]. epiTOC is the average methylation level across 385 probes whereas the expression-based index is the average expression level across 9 genes: *CDKN3, ILF2, KDELR2, RFC4, TOP2A, MCM3, KPNA2, CKS2*, and *CDC2*. For each index, we computed its Spearman correlation with each of the top 5 PCs for each cancer type and called statistical significance at q<0.05. To remove epiTOC-correlated probes, we removed any probes correlated with epiTOC (q<0.05; Pearson correlation) and recomputed methylation PC-driver gene associations for each cancer type. Finally, we used Wilcoxon rank sum test to test the associations between mutated driver genes and each of the two indices and called statistical significance at q<0.05 for each cancer type.

### Differential expression analysis of thyroid carcinoma molecular subtypes

First, we visually identified two THCA molecular subtypes based on driver gene mutations and DNA methylation patterns (Fig 4A): (1) *BRAF*-mutated tumors (the *BRAF* group) and (2) *NRAS*- and *HRAS*-mutated tumors (the *NRAS-HRAS* group). Next, we assembled four differentially expressed gene sets (up- or downregulated in one group but identical or regulated in the opposite direction in another group, compared with controls). The search was restricted to genes whose aberrant expression levels coincided with hyper- or hypomethylated probes associated with *BRAF, HRAS*, and *NRAS* mutations (see section below). Using a hypergeometric model, genes in each of the four sets were tested for enrichment by 1,330 canonical pathways collected by MSigDB [55]. The highly transcribed, differential genes were defined by expression levels that are greater than 10 (in log_2_ RSEM) and at least double those of controls.

### Association between driver gene-associated aberrant methylation and aberrant gene expression in thyroid carcinomas

We sought genes whose aberrant expression was correlated with aberrant methylation in the presence of *BRAF, HRAS*, or *NRAS* mutations. First, we identified probes that fell within 1,500 bp of the transcription start site (TSS) or within gene bodies, and whose β values were significantly correlated with expression levels of the corresponding gene. Because a methylated CpG may up- or downregulate gene expression in a context-dependent manner [50, 56, 57], we computed the significance of the Spearman correlation between each individual probe’s methylation and the expression level of its corresponding gene. We considered gene-probe pairs significant at *q* < 0.05. Second, we analyzed the association between β values and *BRAF, HRAS*, and *NRAS* mutation status for all probes using the two-sided Wilcoxon rank sum test. For each driver gene, aberrantly methylated probes (*q* < 0.05) were classified as hyper- or hypomethylated relative to control samples. Third, we integrated the results from the first and second steps to identify aberrantly methylated probes whose methylation levels were significantly correlated with the expression levels of their corresponding genes for each group of *BRAF*-, *HRAS*-, and *NRAS*-mutated samples. Finally, for each group of mutated samples, significantly up- or downregulated genes were identified from the aberrant methylation-matched genes identified in the third step using the two-sided Wilcoxon rank sum test relative to the expression levels of control samples (*q* < 0.05). For example, when we looked for genes that were upregulated in *BRAF*-mutated samples but exhibited no change or were downregulated in *HRAS*- and *NRAS*-mutated samples, we restricted the search to genes that were hyper- (or hypo-) methylated in *BRAF*-mutated samples but exhibited no change or were hypo- (or hyper-) methylated in *HRAS*- and *NRAS*-mutated samples in the third step. Thus, for each driver gene, we obtained a list of aberrantly methylated probes associated with genes that were aberrantly expressed.

## Acknowledgements

We thank Kristin Harper for editorial assistance.

## Supporting information

**S1 Fig.**
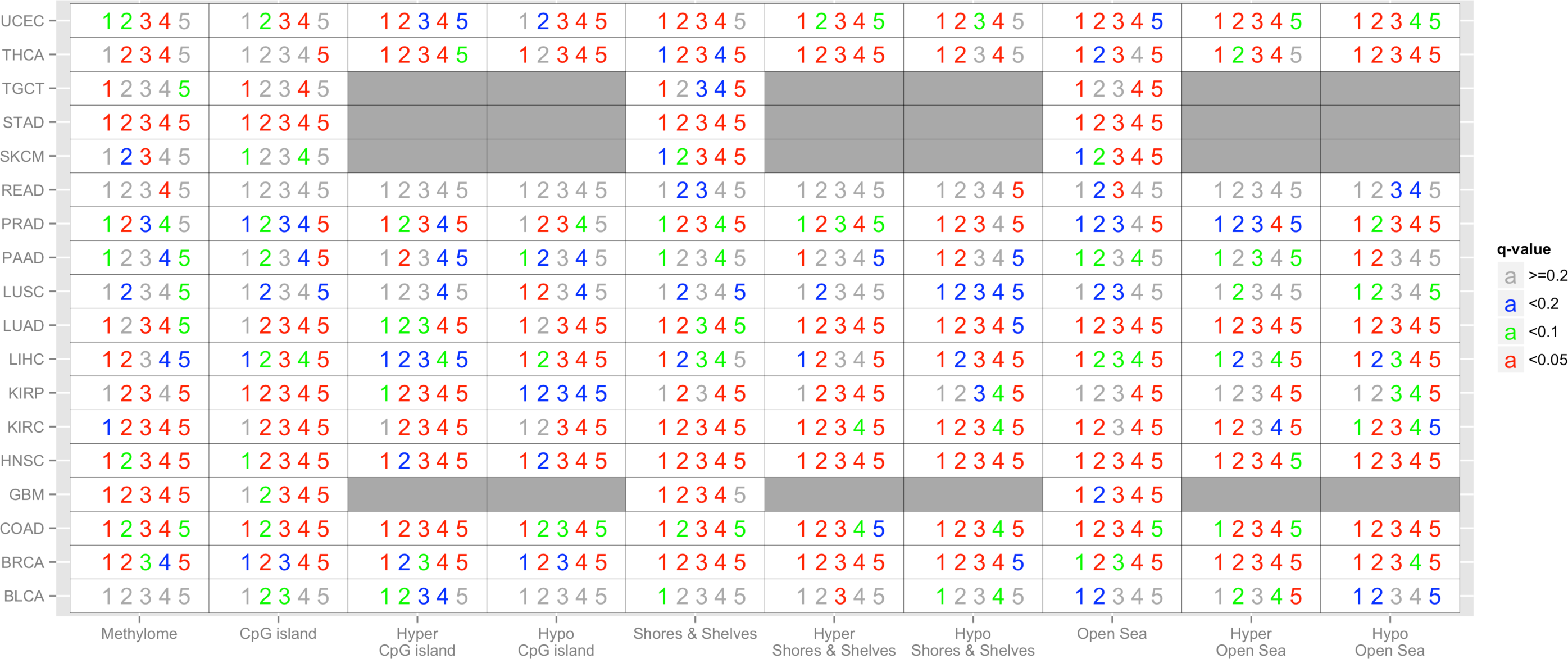
Driver gene-methylation associations across 18 cancer types (*rows*), stratified by CpG subsets (*columns*). The numbers (1–5) indicate the top five principal components (PCs) for each probe set, whereas the color shows the significance of the strongest association between each methylation PC and any driver gene. The probe sets are methylome (all probes), CpG island (CGI) probes, shore and shelf (SS) probes, and open sea probes, further stratified by hyper- and hypomethylation status. For GBM, STAD, SKCM, and TGCT, there were not enough control samples to compute associations for hyper-/hypomethylated probes (shown in dark grey).

**S2 Fig.**
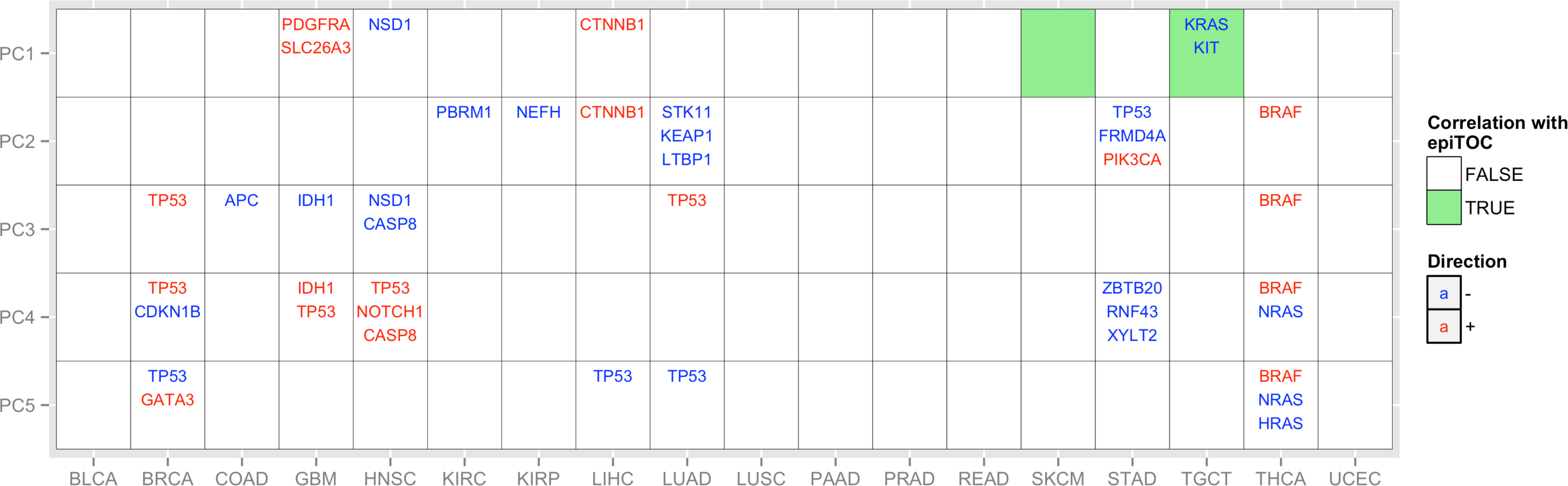
Driver gene mutations are significantly associated with epiTOC-uncorrelated DNA methylation variation in various cancers. The DNAm-based mitotic index called epiTOC (epigenetic Timer Of Cancer) was used to approximate cell proliferation rate in cancer [7]. In 10 of 18 cancer types examined, driver gene mutations were associated with one or more of the top five epiTOC-uncorrelated methylation principal components (PCs). Shown is a grid depicting the three driver genes most significantly associated with each PC. A gene name in blue indicates that mutations in a given gene were significantly associated with the negative extreme of the PC, whereas red indicates association with the positive extreme of the PC. For each PC, the light green background color indicates its correlation with epiTOC (q<0.05; Spearman correlation). See Table 1 for cancer type abbreviation.

**S3 Fig.**
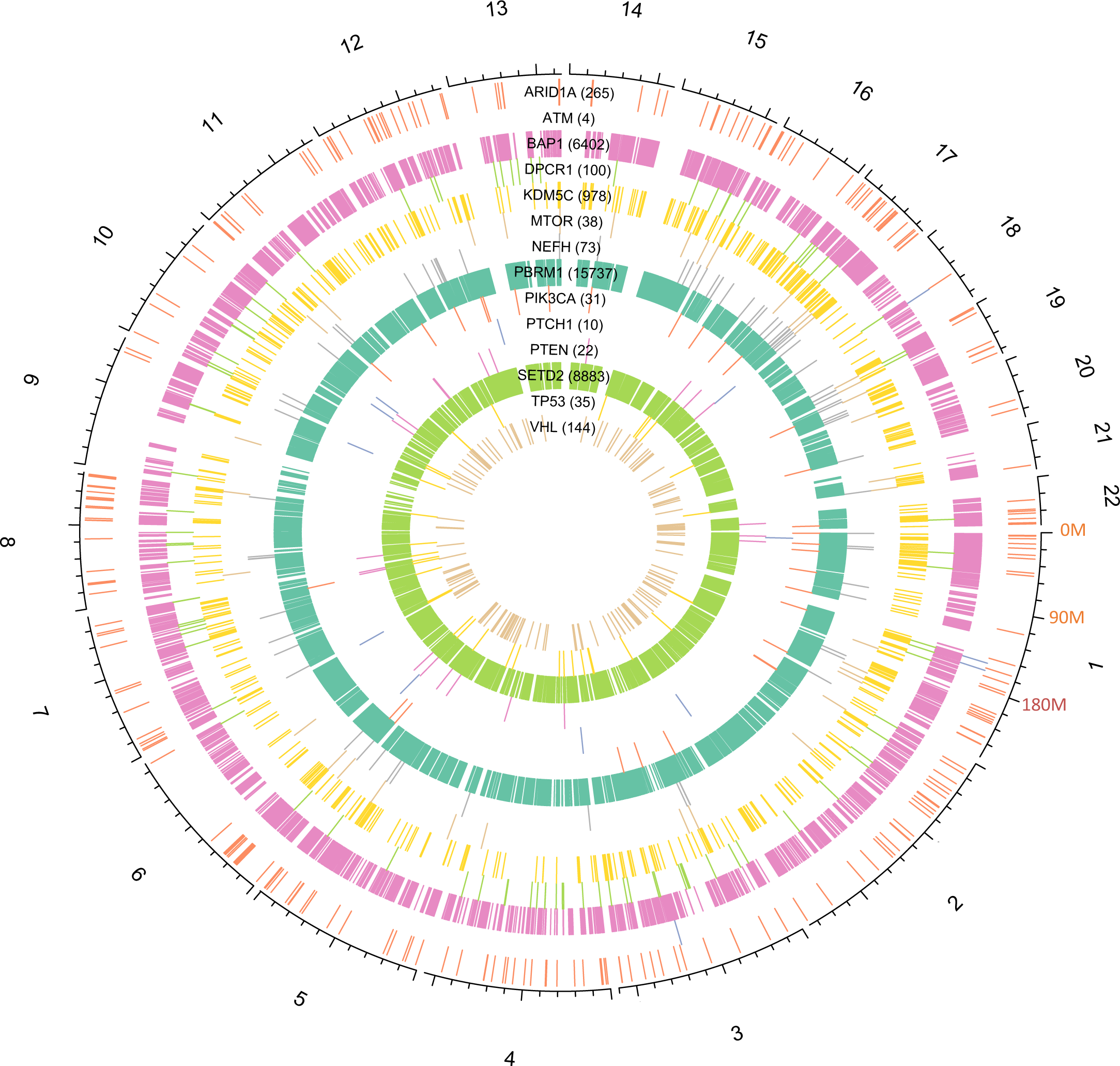
Distribution of driver gene-associated methylation probes throughout the genome in KIRC. Chromosomes 1 to 22 are plotted on a circle, with each chromosome plotted proportional to chromosome length and labeled in the outermost track. The 14 inner tracks correspond to all 14 driver genes in KIRC; gene names and the number of associated probes for each are shown. For each driver gene, associated probes are plotted as line segments in the corresponding track at the appropriate chromosome location. The chromosome length scale is labeled for chromosome 1 (a major interval indicates 90 Mb).

**S4 Fig.**
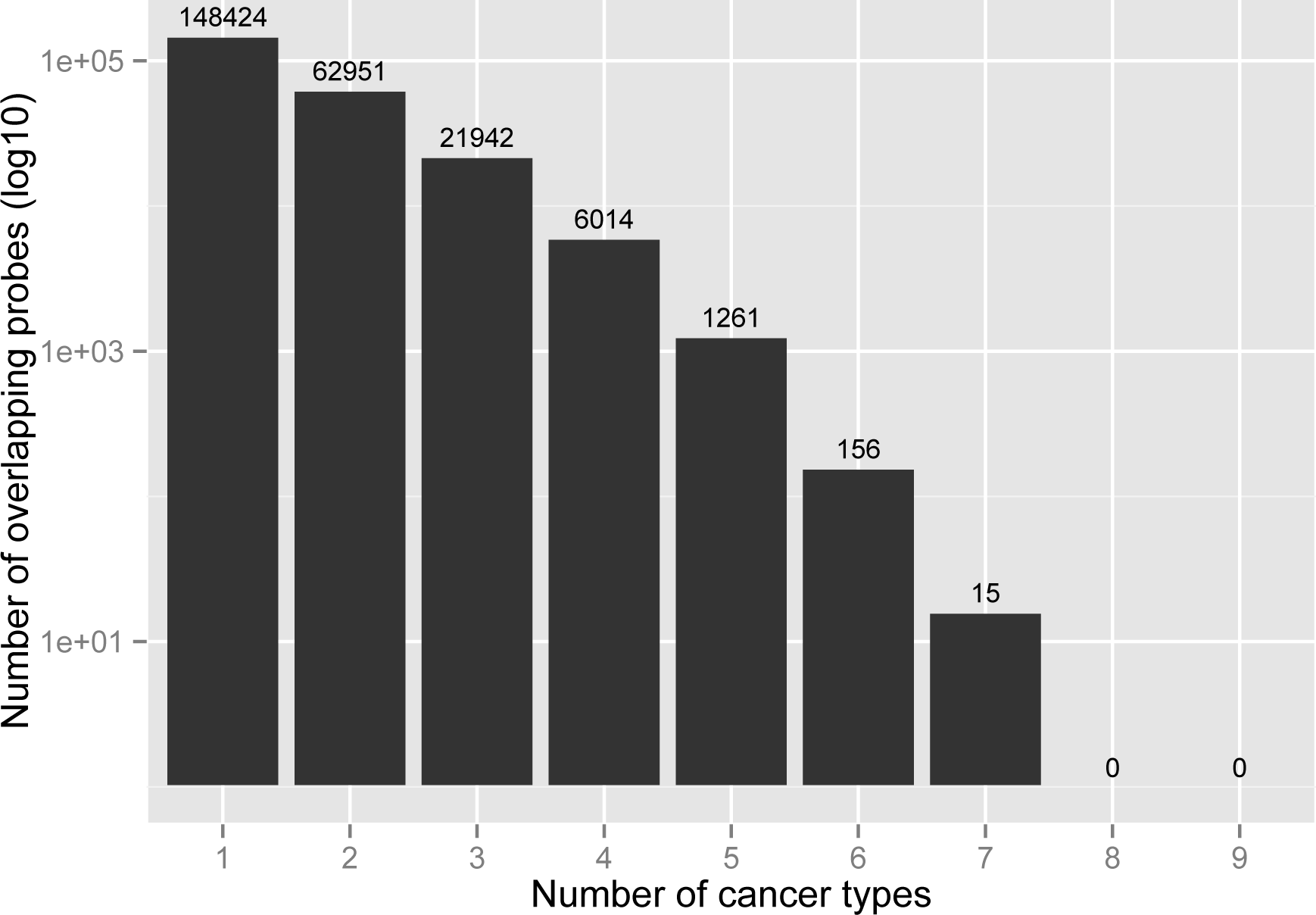
Probes that are negatively associated with *TP53* are shared across cancer types. A bar plot shows the number of *TP53*-negatively associated probes (y-axis in log 10 scale; the number also indicated at the top of each bar) reported in at least 1-9 cancer types (x-axis).

**S5 Fig.**
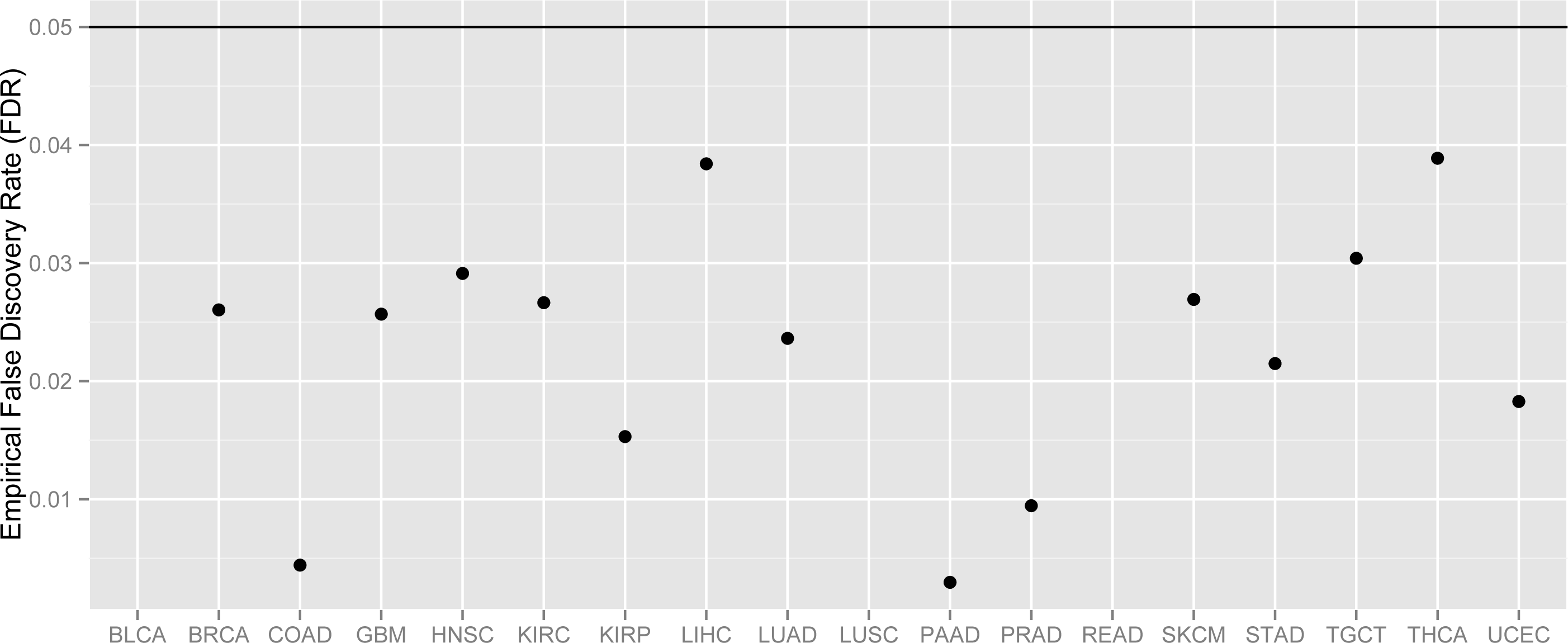
Empirical false discovery rates (FDRs) are controlled by theoretical cutoffs at *q*=0.05 for site-specific associations. The site-specific association was tested between every driver gene and every probe. Significant associations were called at the theoretical FDR (q-value) less than 0.05 for each cancer type. The empirical FDR (y-axis) was estimated for the theoretical cutoff (q=0.05) using the permutation test for each cancer type (*column*). Here, all points are below the line (empirical FDR=0.05), indicating that empirical FDRs are controlled by the theoretical cutoffs.

**S1 Table.** Driver genes associated with the HyperZ and HypoZ indices across 18 cancer types, stratified by direction of association

**S2 Table.** Counts of driver gene-methylation probe associations for 18 cancer types, stratified by CpG subsets: methylome (all probes), CpG island probes, shores and shelves probes, and open sea probes, as well as by hypo- and hypermethylated status (+: positive associations, -: negative associations)

**S3 Table.** Sorted median expression levels of genes differentially expressed (relative to control samples) in the two thyroid carcinoma molecular subtypes identified in Figure 4A: *BRAF*-mutated versus *NRAS*-/*HRAS*-mutated; each Excel tab separates genes by the direction of expression dysregulation (*up* or *down*) and the group of mutated samples showing the expression dysregulation (*BRAF* or *HRAS*/*NRAS*)

**S4 Table.** Pathways enriched with genes dysregulated in the two thyroid carcinoma molecular subtypes identified in Figure 4A: *BRAF*-mutated versus *HRAS-*/*NRAS*-mutated, stratified by the direction of expression dysregulation (*up* or *down*) and the mutant group (*BRAF* or *HRAS*/*NRAS*)

